# Biochemical characterization of RAD52-mediated D-loop formation using fluorophore-labeled DNA substrates

**DOI:** 10.1101/2022.02.23.481227

**Authors:** Kazuki Kamoi, Mika Saotome, Chiaki Kinoshita, Ryohei Tsuchiya, Wataru Kagawa

## Abstract

The human RAD52 protein is thought to have multiple roles in the mechanisms of repairing DNA double-strand breaks that are caused by replication errors and reactive oxygen species. One such role is to mediate the formation of a displacement loop (D-loop), which is a critical reaction intermediate in homologous recombinational repair. RAD52 is suggested to promote the formation of D-loops when facilitating DNA synthesis at stalled or collapsed replication forks during mitosis. However, RAD52-mediated D-loop formation remains poorly characterized, and the detailed molecular mechanism of the D-loop formation reaction catalyzed by RAD52 is still unclear. In the present study, we developed a gel-based assay that enables rapid detection of RAD52-mediated D-loop formation. This assay utilizes a fluorophore-labeled, single-stranded DNA substrate. In addition to the rapid detection of D-loops, D-loop extension was observed when DNA polymerase was added to the reaction. This assay can also be used for screening large numbers of compounds that either stimulate or inhibit RAD52-mediated D-loop formation. The D-loop formation assay developed in this study is potentially useful for mechanistic studies of DSB repair involving RAD52-mediated D-loop formation, as well as for screening compounds with potential therapeutic effects.

## Introduction

Genomic DNA is constantly damaged by replication errors and reactive oxygen species that are produced as byproducts of normal metabolism of oxygen. These endogenous factors often cause DNA double-strand breaks (DSBs), which pose a threat to genome integrity. Failure to properly repair DSBs may cause cell death, or induce genomic rearrangements that can lead to cancer [1]. Homologous recombination (HR) is a widely conserved mechanism for the accurate repair of DSBs. The RAD52 protein is believed to be one of the key repair factors functioning in the eukaryotic HR [2]. RAD52 display a multitude of DNA recombination activities, such as single-strand DNA annealing, strand exchange, and D-loop formation, all of which are likely to be important in the various HR pathways [3].

Much focus has been devoted to the single-strand DNA annealing activity of RAD52. RAD52 plays a critical role in single-strand annealing (SSA), a DSB repair pathway that takes place between tandem repeats [4]. Both yeast and human RAD52 proteins catalyze the annealing reaction between homologous single-stranded DNAs (ssDNA) *in vitro* [5-7]. RAD52 is capable of promoting the annealing reaction between replication protein A(RPA)-coated ssDNA, which suggests the functional association between RAD52 and RPA during SSA. X-ray crystallographic studies of the RAD52-ssDNA complexes have revealed the molecular details of the interaction between RAD52 and ssDNA [8], and suggest an annealing mechanism in which complementary DNA strands are trapped between two RAD52 rings [8,9].

Increasing evidences suggest that RAD52 promotes D-loop formation *in vivo*. In yeast cells lacking a functional RAD51 recombinase, RAD52 is likely the protein that promotes break-induced replication (BIR), which involves the formation of D-loops between the broken chromosome and the undamaged chromosome [10]. BIR has been reported to occur in higher eukaryotes as well, in cells experiencing DNA replication stress. In these cells, unscheduled DNA synthesis was shown to occur in mitosis through BIR, and it is reported to be dependent on RAD52 [11,12]. RAD52-dependent BIR has also been suggested to occur during a telomerase-independent, telomere lengthening mechanism called, alternative lengthening of telomeres (ALT) [13]. Because a significant fraction of cancer cells relies on ALT to maintain telomeres, ALT is a potential target for drug development.

Despite the importance of RAD52-mediated D-loop formation in multiple DSB repair pathways, its mechanism remains largely unclear. A better understanding of the molecular mechanisms underlying RAD52-mediated D-loop formation will greatly facilitate the characterizations of the different DSB repair pathways in which RAD52 is involved. In the present study, we developed an assay to rapidly assess the D-loop formation activity of RAD52. Using this assay, we found that both D-loop formation and DNA synthesis can be monitored in the same reaction. This assay is potentially useful for future investigations to dissect the molecular events in RAD52-dependent DSB repair.

## Materials and Methods

### Proteins

The full-length, human RAD52 protein was overexpressed in the *E. coli* strain JM109(DE3). The strain was co-transformed with a pET-15b vector containing the human RAD52 gene and the pArg3Arg4 vector [14]. For a typical preparation, the co-transformed *E. coli* strain was cultured in 800 mL of LB medium at 30°C. When the culture reached an optical density (A600) of ∼0.6, RAD52 expression was induced with 0.5 mM isopropyl 1-thio-β-D-galactopyranoside (final concentration) for ∼16 hours. Afterward, the cells were harvested, resuspended in ∼40 mL of ice-cold, Lysis Buffer (50 mM Tris-HCl, pH 7.8, 0.3 M KCl, 10% glycerol, 10 mM imidazole), and lysed by sonication. The cell lysate was cleared of insoluble material by centrifugation at 35,200 x *g* for 30 min. The resulting supernatant was mixed with 2 mL of Ni-NTA agarose beads, and incubated with gentle mixing for 1 h. The RAD52-bound, Ni-NTA agarose beads were packed into an Econo-column, and washed with 90 mL of Wash Buffer (50 mM Tris-HCl, pH 7.8, 0.3 M KCl, 10% glycerol, 50 mM imidazole). RAD52 was eluted with a 90-mL linear gradient of 50-400 mM imidazole. Peak fractions were combined, and dialyzed against RAD52 Buffer (20 mM HEPES-KOH, pH 7.5, 0.5 mM EDTA, 2 mM 2-mercaptoethanol, 5% glycerol, 0.2 M KCl). Prior to dialysis, 3 units of thrombin protease per mg of RAD52 were added to cleave off the hexahistidine tag. The dialyzed RAD52 fraction was loaded onto a 3 mL SP-Sepharose column. After washing the column with 60 mL of RAD52 Buffer, the protein was eluted with a 60-mL linear gradient of 0.2–0.8M KCl. Peak fractions were collected, filtered through a 5-μm syringe filter, and concentrated to approximately 2 mg/mL using an Vivaspin Turbo 15 centrifugal filter (100K MWCO). The concentrated RAD52 was aliquoted in small volumes (50 μL), flash frozen in liquid nitrogen, and stored at -80°C. RAD52 concentration was determined by measuring the absorption at 280 nm, using extinction coefficients calculated with the Protparam tool on the ExPASy website (http://web.expasy.org/protparam/).

### DNA substrates

For the D-loop formation assay, an HPLC-grade, 60-mer oligonucleotide with a cyanine dye 5 (Cy5) covalently attached to the 5’ terminus, and a negatively supercoiled, plasmid DNA (pUC18, 2,680 base pairs) were used as ssDNA and dsDNA substrates, respectively. The oligonucleotide is homologous to the pUC18 DNA sequence 1,501-1,560 (5’-GAA AAC TCA CGT TAA GGG ATT TTG GTC ATG AGA TTA TCA AAA AGG ATC TTC ACC TAG ATC-3’). The pUC18 plasmid DNA was purified from sarkosyl-lysed cells, as described previously [15]. The plasmid DNA was purified by agarose gel electrophoresis using the Model 491 Prep Cell (Bio-Rad). All DNA concentrations are expressed in moles of molecules.

### D-loop formation assay

D-loops were formed in a 10-μL reaction mixture containing 83.3 nM ssDNA, 18.7 nM pUC18, a 2-μL aliquot of RAD52, and a 2-μL aliquot of a 5-fold reaction mix (0.1 M Hepes-KOH, pH 7.5, 10 mM 2-mercaptoethanol). For a typical reaction, ssDNA was pre-incubated with RAD52 at 37°C for 5 min. Afterward, pUC18 was added, and the mixture was further incubated for 15 min. D-loops were deproteinized by adding 2 μL of 5% lithium dodecyl sulfate and 1 μL of 20 mg/mL proteinase K (New England Biolabs), and incubating the mixture at 37°C for 15 min. The resulting reaction mixture (13 μL) was thoroughly mixed with 5 μL of 40% sucrose, and fractionated through a 1% agarose gel in 1x TAE buffer for 2 hours at 3.3 V/cm. The gels were visualized using a Typhoon FLA 7000 image scanner (Cytiva). Bands were quantified using the ImageJ software [16]. Because the ssDNA was in excess compared with the dsDNA, the yield of D-loops was expressed as the percentage of pUC18 dsDNA incorporated into D-loops. For the reaction containing various compounds, compounds included in the Additive Screening Kit (Hampton Research) were used.

### D-loop extension assay

D-loops were formed as described above. Following the 15 min-incubation after the addition of pUC18, 1 μL of 2 mM dNTP and 0.5 μL (2.5 U) of DNA polymerase I Klenow fragment (New England Biolabs) were added to the reaction mixture, and incubated for indicated times. Afterward, products were deproteinized (as described above), and heated at 90°C for 1 min, followed by quick chilling on ice water. The resulting reaction mixture (14.5 μL) was thoroughly mixed with 5 μL of 40% sucrose, and fractionated through a 12% polyacrylamide gel in 0.5x TBE buffer. The gels were visualized using a Typhoon FLA 7000 image scanner.

## Results

### RAD52-mediated D-loop formation detected using a fluorophore-labeled ssDNA substrate

Previously, we and others have used gel assays to show that RAD52 mediates the formation of D-loops, which are key reaction intermediates in homologous recombinational repair [14,17-19]. In those assays, ssDNA substrates were labeled with ^32^P to detect D-loops. Radioisotope labeling is often the choice for labeling oligonucleotides, because it allows highly sensitive detection of products. A drawback of using radiolabeled DNA substrates is that the detection is time consuming. Gels containing the fractionated, radiolabeled substrates are dried, and immobilized on an ion exchange paper prior to visualization. Furthermore, DNA substrates labeled with radioisotopes are not suited for long-term storage, and frequent radiolabeling may be required to minimize signal strength differences between different experiments. Fluorophore labeling, on the other hand, can overcome these drawbacks associated with radioisotope labeling. Fluorophore-labeled DNA can be visualized immediately after electrophoresis with an image scanner like Typhoon (Cytiva). They can be stored for long periods of time at low temperatures (e.g. -30°C), protected from light.

To examine whether a fluorophore-labeled ssDNA can substitute for ^32^P-labeled ssDNA in the RAD52-mediated D-loop formation assay, we used a 60-mer oligonucleotide with a Cyanine dye 5 (Cy5) conjugated to the 5’ terminus (Fig. 1A). Consistent with our previous studies [14,20], RAD52 exhibited a concentration-dependent, D-loop formation activity (Fig. 1B and C). Maximum activity was observed at 1.5 μM RAD52, and the activity sharply dropped at slightly higher or lower RAD52 concentrations. Hence, the D-loop formation activity was detected within a narrow range of RAD52 concentrations. Also consistent with our previous studies, when RAD52 was preincubated with the dsDNA substrate, followed by the addition of the ssDNA substrate, D-loop formation was significantly diminished (Fig. 1D). Thus, when RAD52 associates with dsDNA before ssDNA, it becomes incompatible for promoting the formation of D-loops.

**Fig. 1.**
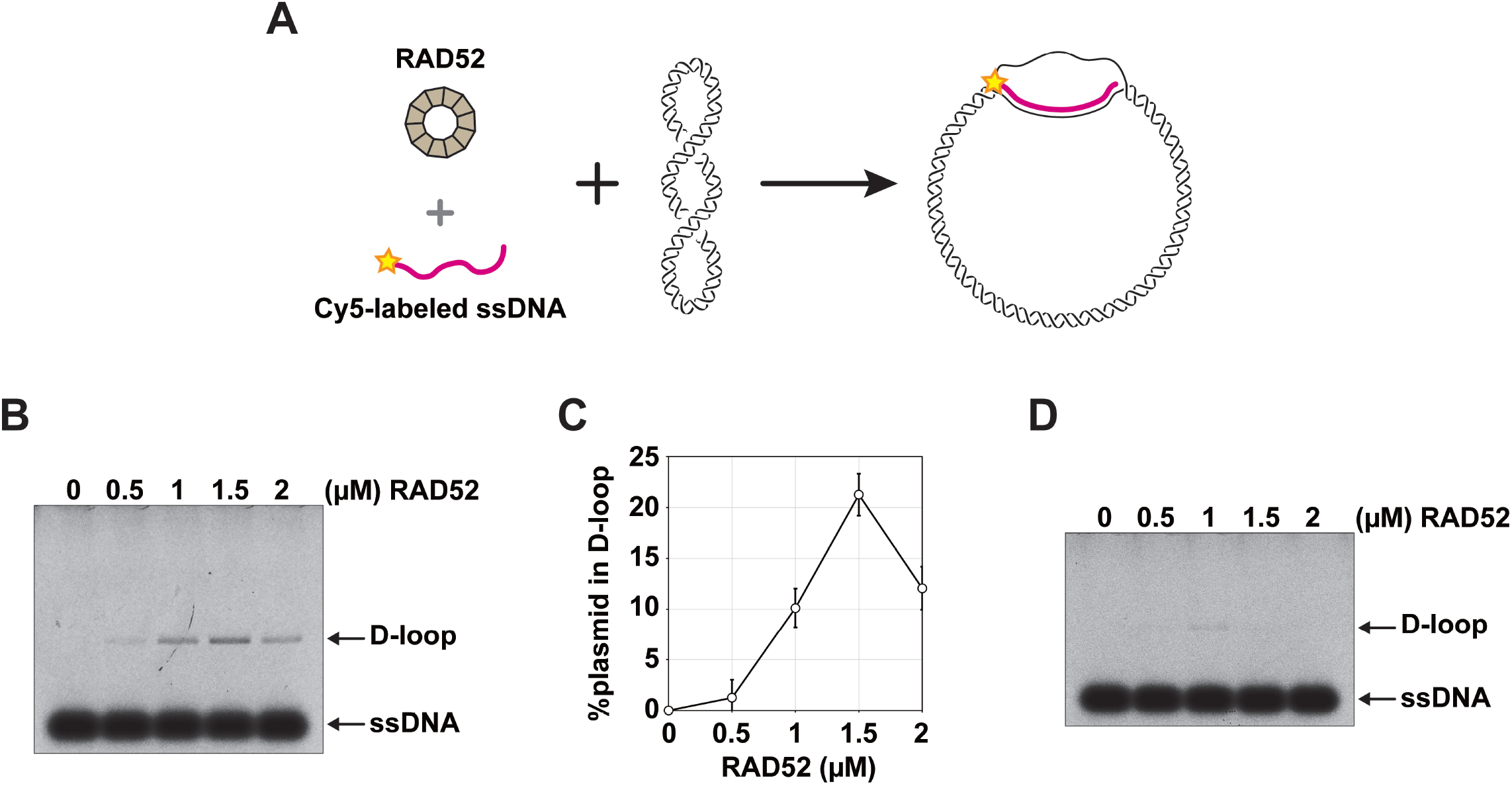
A gel-based assay to detect RAD52-mediated D-loop formation using fluorophore-labeled DNA substrates. **A**, Schematic diagram of the D-loop formation reaction. A 60-mer oligonucleotide labeled with Cyanine 5 at its 5’ end was used as the ssDNA substrate. For the dsDNA substrate, a negatively supercoiled, pUC18 (2,680 bp) was used. **B**, D-loop formation catalyzed by the human RAD52 protein. The ssDNA substrate (83 nM) was pre-incubated with RAD52, prior to the addition of dsDNA (18.7 nM). Reaction solutions were fractionated through an 1% agarose gel. **C**, Graphical representation of the amount of D-loops as a function of RAD52 concentration. **D**, Inverse D-loop formation reaction, in which the dsDNA substrate (18.7 nM) was initially pre-incubated with RAD52, followed by the addition of the ssDNA substrate (83 nM).

### Visualization of D-loop extension

We next investigated whether the D-loop formation assay utilizing the fluorophore-labeled ssDNA can be used for detecting the extension of D-loops by DNA polymerases. To do so, we added dNTPs and the DNA polymerase I (Klenow fragment) to the D-loop reaction mixture, followed by deproteinization and heat treatment to dissociate the D-loops (Fig. 2A). The resulting reaction mixture was fractionated through a polyacrylamide gel, and the Cy5-labeled ssDNA was immediately visualized by an image scanner (Typhoon). Multiple bands with migration distances that were clearly shorter than that of the unextended, Cy5-labeled ssDNA were observed (Fig. 2B). These results suggest that the D-loop formation assay developed in this study can be used for monitoring the D-loop extension reaction in the presence of RAD52 and DNA polymerases.

**Fig. 2.**
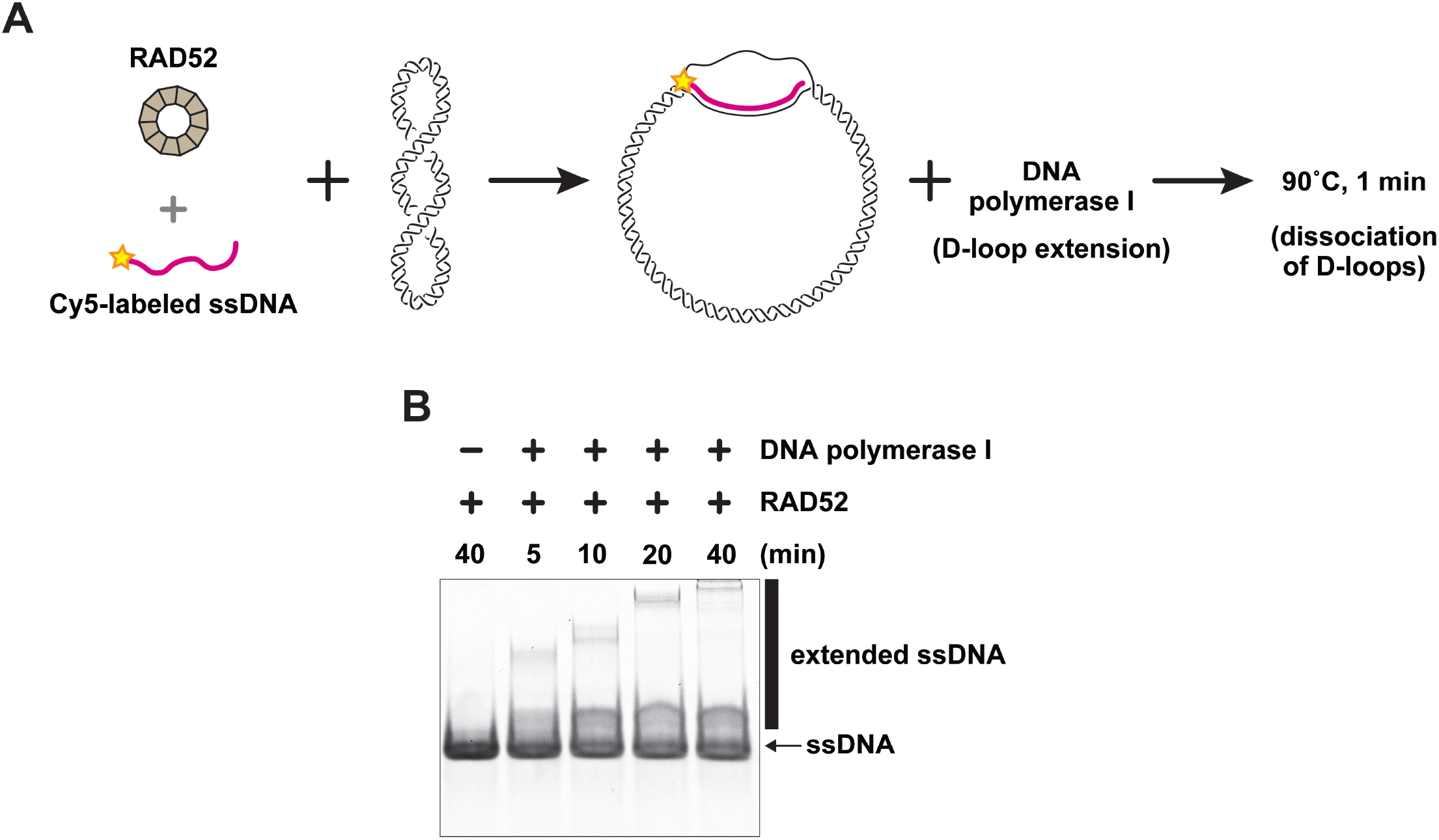
D-loop extension by the DNA polymerase I (Klenow fragment). **A**, Schematic diagram of the D-loop formation and D-loop extension reactions. **B**, D-loop extension catalyzed by the DNA polymerase I (Klenow fragment). Prior to the addition of the DNA polymerase I (Klenow fragment) to the reaction mixtures, D-loops were formed in the presence of RAD52 (1.5 μM). Reaction solutions were fractionated through a 12% polyacrylamide gel.

### Rapid screening for compounds that affect RAD52-mediated D-loop formation

The D-loop formation assay using a Cy5-labeled ssDNA yielded similar results compared to the assay using ^32^P-labeled ssDNA established in our previous studies. The assay utilizing a fluorophore-labeled ssDNA can be completed in about half the time compared with the assay using the ^32^P-labeled ssDNA. This enabled us to rapidly screen for compounds that stimulate or inhibit RAD52-mediated D-loop formation. We tested a total of 93 compounds (Table 1), by separately adding them to a D-loop reaction mixture in the presence of 1.5 μM RAD52 (Fig. 3). D-loop formation was stimulated in the presence of 1 mM calcium, magnesium, and manganese ions (Fig. 3, compound numbers 3, 6, 7). To further examine the effects of these divalent cations, RAD52-mediated D-loop formation was monitored at several concentrations of magnesium chloride and calcium chloride between 0 and 10 mM (Fig. 4A and B). Both magnesium and calcium ions had the strongest stimulatory effect at 1 mM. The yields of D-loops were approximately 2-fold of that without the addition of these ions. Concentrations higher than 1 mM of these divalent ions were inhibitory towards D-loop formation. We also examined the effect of potassium ion, because all D-loop reaction mixtures, by default, contained 20 mM potassium chloride that was carried over from the RAD52 protein solution. Increasing the concentration of potassium ions resulted in the pronounced decrease in D-loop formation (Fig. 4C).

**Table 1.** List of compounds tested for their effects on RAD52-mediated D-loop formation.

|  | type of reagent | compound name | final concentration | effect |
| --- | --- | --- | --- | --- |
| 1 | multivalent ion | barium chloride | 1 mM |  |
| 2 | multivalent ion | cadmium chloride | 1 mM |  |
| 3 | multivalent ion | calcium chloride | 1 mM | + |
| 4 | multivalent ion | cobalt(II) chloride | 1 mM | - |
| 5 | multivalent ion | copper(II) chloride | 1 mM | - |
| 6 | multivalent ion | magnesium chloride | 1 mM | + |
| 7 | multivalent ion | manganese(II) chloride | 1 mM | + |
| 8 | multivalent ion | strontium chloride | 1 mM |  |
| 9 | multivalent ion | yttrium(III) chloride | 1 mM | - |
| 10 | multivalent ion | zinc chloride | 1 mM | - |
| 11 | multivalent ion | iron(III) chloride | 1 mM |  |
| 12 | multivalent ion | nickel(II) chloride | 1 mM | - |
| 13 | multivalent ion | chromium(III) chloride | 1 mM |  |
| 14 | multivalent ion | praseodymium(III) acetate | 1 mM |  |
| 15 | salt | ammonium sulfate | 10 mM |  |
| 16 | salt | potassium chloride | 10 mM |  |
| 17 | salt | lithium chloride | 10 mM |  |
| 18 | salt | sodium chloride | 20 mM |  |
| 19 | salt | sodium fluoride | 5 mM |  |
| 20 | salt | sodium iodide | 10 mM |  |
| 21 | salt | sodium thiocyanate | 20 mM |  |
| 22 | salt | potassium sodium tartrate | 10 mM |  |
| 23 | salt | sodium citrate | 10 mM |  |
| 24 | salt | cesium chloride | 10 mM |  |
| 25 | salt | sodium malonate, pH 7.0 | 10 mM |  |
| 26 | amino acid | L-proline | 1 mM |  |
| 27 | dissociating agent | phenol | 1 mM |  |
| 28 | dissociating agent | dimethyl sulfoxide | 0.3 % v/v |  |
| 29 | dissociating agent | sodium bromide | 1 mM |  |
| 30 | linker | 6-aminohexanoic acid | 0.3 % w/v |  |
| 31 | linker | 1,5-diaminopentane | 0.3 % w/v | - |
| 32 | linker | 1,6-diaminohexane | 0.3 % w/v | - |
| 33 | linker | 1,8-diaminooctane | 0.3 % w/v | - |
| 34 | linker | glycine | 10 mM |  |
| 35 | linker | taurine | 1 mM |  |
| 36 | linker | betaine | 1 mM |  |
| 37 | polyamine | spermidine | 1 mM | - |
| 38 | polyamine | spermine | 1 mM | - |
| 39 | polyamine | hexamine cobalt(III) chloride | 1 mM | - |
| 40 | polyamine | sarcosine | 1 mM |  |
| 41 | chaotrope | trimethylamine | 1 mM |  |
| 42 | chaotrope | guanidine hydrochloride | 10 mM |  |
| 43 | chaotrope | urea | 1 mM |  |
| 44 | co-factor | β-nicotinamide adenine dinucleotide | 1 mM | |
| 45 | co-factor | adenosine-5'-triphosphate | 1 mM |  |
| 46 | reducing agent | TCEP | 1 mM |  |
| 47 | reducing agent | L-glutathione | 0.1 mM |  |
| 48 | polymer | polyvinylpyrrolidone K15 | 0.05 % w/v |  |
| 49 | polymer | dextran sulfate | 0.3 % w/v | – |
| 50 | polymer | pentaerythritol ethoxylate (3/4 EO/OH) | 0.4 % v/v |  |
| 51 | polymer | polyethylene glycol 3,350 | 0.1 % w/v |  |
| 52 | carbohydrate | sucrose | 0.3 % w/v |  |
| 53 | carbohydrate | xylitol | 0.3 % w/v |  |
| 54 | carbohydrate | D-sorbitol | 0.3 % w/v |  |
| 55 | carbohydrate | myo-inositol | 0.12 % w/v |  |
| 56 | carbohydrate | D-(+)-trehalose | 0.3 % w/v |  |
| 57 | carbohydrate | D-(+)-galactose | 0.3 % w/v |  |
| 58 | polyol | ethylene glycol | 0.3 % v/v |  |
| 59 | polyol | glycerol | 0.3 % v/v |  |
| 60 | non-detergent sulfobetaine | NDSB-195 | 30 mM |  |
| 61 | non-detergent sulfobetaine | NDSB-201 | 20 mM |  |
| 62 | non-detergent sulfobetaine | NDSB-211 | 20 mM |  |
| 63 | non-detergent sulfobetaine | NDSB-221 | 20 mM |  |
| 64 | non-detergent sulfobetaine | NDSB-256 | 10 mM |  |
| 65 | amphiphile | CYMAL® -7 | 1.5 µM |  |
| 66 | amphiphile | benzamidine | 0.2 % w/v | – |
| 67 | detergent | n-dodecyl-N,N-dimethylamine-N-oxide | 0.05 % w/v |  |
| 68 | detergent | n-octyl-β-D-glucoside | 0.05 % w/v |  |
| 69 | osmolyte | n-dodecyl-β-D-maltoside | 0.05 % w/v |  |
| 70 | organic solvent | trimethylamine N-oxide | 0.3 % w/v |  |
| 71 | organic solvent | 1,6-hexanediol | 0.3 % w/v |  |
| 72 | organic solvent | (+/-)-2-methyl-2,4-pentanediol | 0.3 % v/v |  |
| 73 | organic solvent | polyethylene glycol 400 | 0.5 % v/v |  |
| 74 | organic solvent | Jeffamine M-600 pH 7.0 | 0.5 % v/v |  |
| 75 | organic solvent | 2,5-hexanediol | 0.4 % v/v |  |
| 76 | organic solvent | (±)-1,3-butanediol | 0.4 % v/v |  |
| 77 | organic solvent | polypropylene glycol P 400 | 0.4 % v/v |  |
| 78 | organic solvent | 1,4-dioxane | 0.3 % v/v |  |
| 79 | organic solvent | ethanol | 0.3 % v/v |  |
| 80 | organic solvent | 2-propanol | 0.3 % v/v |  |
| 81 | organic solvent | methanol | 0.3 % v/v |  |
| 82 | organic solvent | 1,2-butanediol | 0.1 % v/v |  |
| 83 | organic solvent | tert-butanol | 0.4 % v/v |  |
| 84 | organic solvent | 1,3-propanediol | 0.4 % v/v |  |
| 85 | organic solvent | acetonitrile | 0.4 % v/v |  |
| 86 | organic solvent | formamide | 0.4 % v/v |  |
| 87 | organic solvent | 1-propanol | 0.4 % v/v |  |
| 88 | organic solvent | ethyl acetate | 0.05 % v/v |  |
| 89 | organic solvent | acetone | 0.4 % v/v |  |
| 90 | organic solvent | dichloromethane | 0.0025 % v/v |  |
| 91 | organic solvent | 1-butanol | 0.07 % v/v |  |
| 92 | organic solvent | 2,2,2-trifluoroethanol | 0.4 % v/v |  |
| 93 | organic solvent | 1,1,1,3,3,3-hexafluoro-2-propanol | 0.4 % v/v |  |

"+" indicates that the reagent stimulated D-loop formation by RAD52.
"–" indicates that the reagent inhibited D-loop formation by RAD52.

**Fig. 3.**
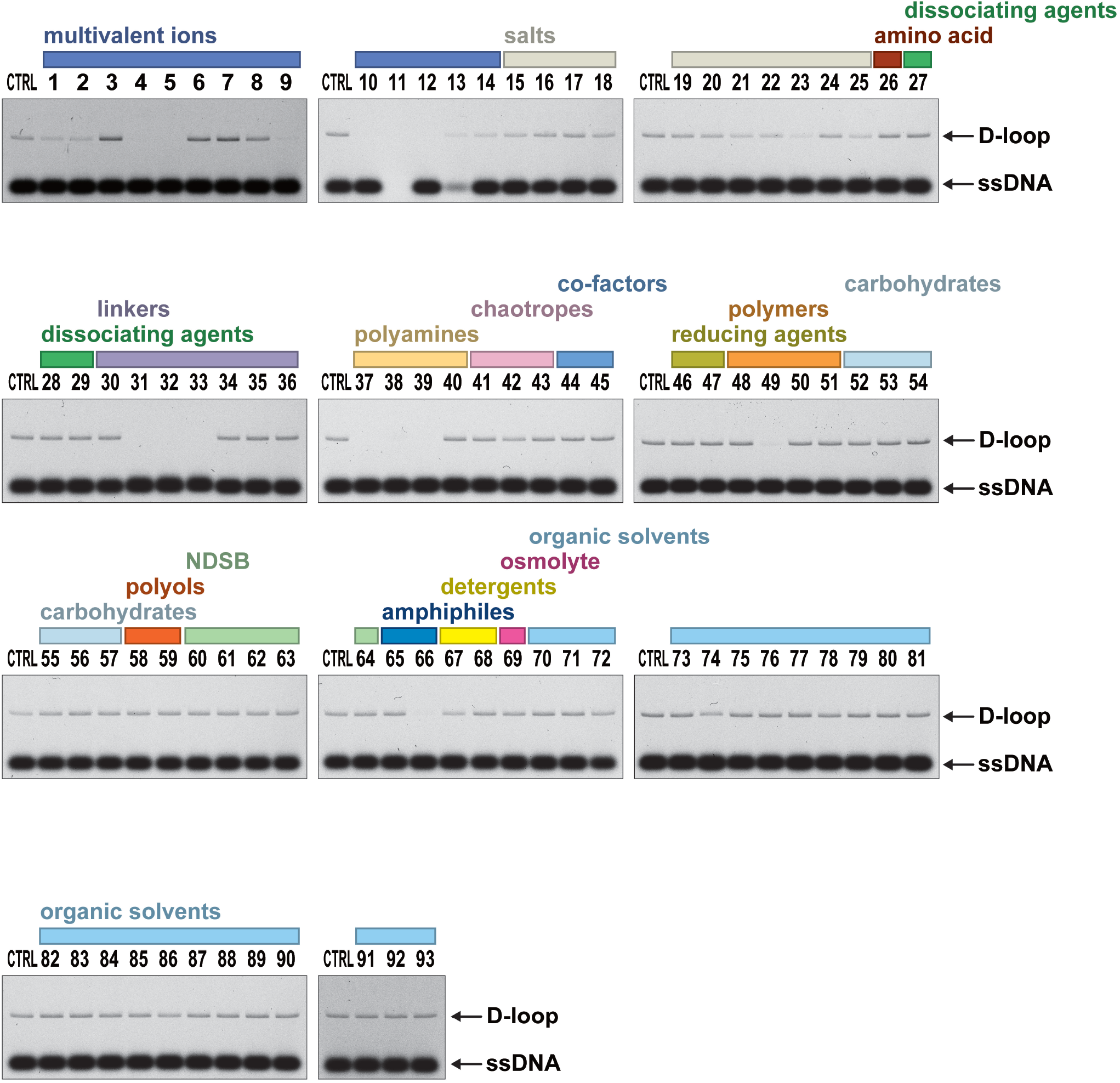
Effects of various compounds on RAD52-mediated D-loop formation. For all reactions, RAD52 (1.5 μM) was pre-incubated with ssDNA (83 nM) in the presence of the indicated compound, followed by the addition of dsDNA (18.7 nM). The D-loop formation reaction without the addition of compounds is indicated as “CTRL”. Final concentrations of the compounds in the reaction mixture are shown in Table 1.

**Fig. 4.**
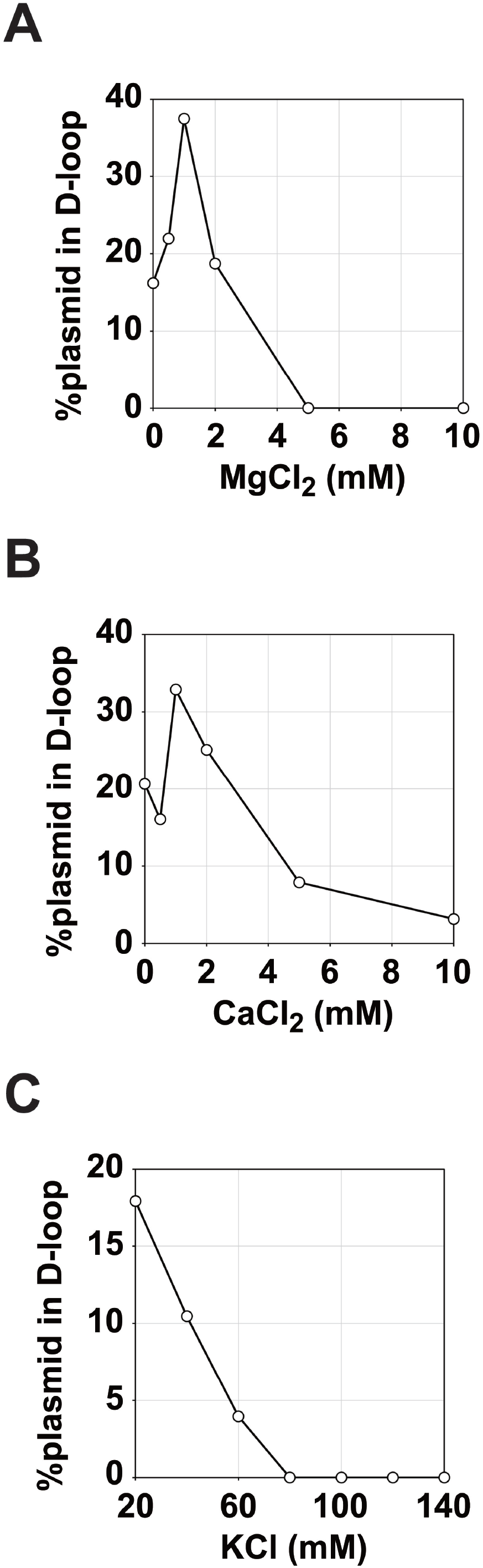
Effects of potassium, magnesium, and calcium ions on RAD52-mediated D-loop formation. In all three reactions (shown in A-C), RAD52 (1.5 μM) was pre-incubated with ssDNA (83 nM) in the presence of the indicated concentrations of magnesium, calcium, or potassium ions, followed by the addition of dsDNA (18.7 nM). Graphical representations of the amount of D-loops as a function of magnesium chloride concentration (**A**), calcium chloride concentration (**B**), and potassium chloride concentration (**C**).

Multiple compounds were found to completely inhibit RAD52-mediated D-loop formation (Fig. 3, compound numbers 4, 5, 9, 10, 12, 31-33, 37-39, 49, 66). Despite the significant number of these compounds, the types of reagents were limited (14 compounds were categorized into 6 groups) (see Table 1). They were divalent (cobalt, copper, zinc, nickel) or trivalent (yttrium) ions, linkers (1,5-diaminopentane, 1,6-diaminohexane, 1,8-diaminooctane), polyamines (spermidine, spermine, hexamine cobalt(III) chloride), polymer (dextran sulfate), and amphiphiles (benzamidine).

## Discussion

In the present study, we found that fluorophore-labeled ssDNA substrates can be used for detecting RAD52-mediated D-loop formation. In cases where fluorescent dye provides sufficient signal for product detection, the use of fluorescent dyes may be advantageous over the use of radioisotopes. The biggest benefit of using fluorescent dyes could be the rapid detection of products. For the D-loop formation assay, agarose gels are used to fractionate the plasmid-sized D-loops. If ssDNA is labeled with radioisotopes (such as ^32^P), the agarose gel must be dried, and the fractionated products must be immobilized on an ion-exchange paper, which takes considerable amount of time. By contrast, if ssDNA is labeled with a fluorescent dye, products can be immediately visualized with an image scanner, after electrophoresis.

We have taken advantage of the developed D-loop formation assay, and rapidly screened for compounds that affect RAD52-mediated D-loop formation. We found that most of the compounds that affected RAD52 activity were either multivalent ions, linkers, or polyamines. Many of these compounds are known to directly interact with DNA. Thus, stimulation or inhibition of D-loop formation by these compounds could be through direct interactions, such as stabilization of reaction intermediates or steric hinderance, or by indirect means, such as inducing a particular DNA structure nearby the D-loop structure. We cannot exclude the possibility that these compounds directly interact with RAD52. Structural studies of RAD52 bound to ssDNA revealed a spherical density in the DNA binding site that is likely accommodated by a metal ion [8]. While much work is needed to decipher the molecular mechanisms underlying the stimulation or inhibition of D-loop formation by these compounds, the developed D-loop formation assay provides a valuable tool for rapidly assessing the effects of various compounds and reaction conditions.

Lastly, we have demonstrated that the D-loop formation assay using fluorophore-labeled ssDNA can be utilized to detect D-loop extension by DNA polymerases. RAD52 has been implicated in directly promoting the formation of D-loops during mitosis [11,12]. RAD52 is also shown to play important roles in alternative-lengthening of telomeres (ALT), possibly mediating the formation of D-loops [13]. In these processes, RAD52 may function in conjunction with DNA polymerases to extend the D-loops, and promote DNA repair. The D-loop formation assay established in the present study may facilitate biochemical studies of RAD52 and DNA polymerases to clarify the precise roles of these proteins in these repair events.

## Author contributions

W.K., M.S., and K.K. designed the research; K.K., M.S., C.K., and R.T. performed the research; K.K., M.S., C.K., R.T., and W.K. analyzed the data; W.K. wrote the manuscript.

## Funding

This study was funded by JSPS KAKENHI Grant Numbers: 16H01316, 19K12328, and 20H00449 (to W.K.). W.K. was also supported by the Science Research Promotion Fund from the Japan Private School Promotion Foundation, and by the Priority Research Funding from Meisei University.

## Conflict of Interest

The authors declare that they have no conflict of interest.

## Notes

### Competing Interest Statement

The authors have declared no competing interest.

